# The VGLUT3-p.T8I Mutation Alters Striatal Acetylcholine Dynamics in Response to Social Stressors and Drives Selective Social Avoidance in Male Mice

**DOI:** 10.64898/2026.09.09.750466

**Authors:** Marine Pujol, Véronique Bernard, Salah El Mestikawy, Véronique Fabre, Stéphanie Daumas

**Author notes:** Correspondence: Veronique Fabre : Cassan, bâtiment B, étage 4, porte 425-E, case courrier 37, 7 quai Saint Bernard, 75 252 PARIS CEDEX;, Stephanie Daumas : Cassan, bâtiment B, étage 4, porte 425-E, case courrier 37, 7 quai Saint Bernard, 75 252 PARIS CEDEX;, Marine Pujol : Perry Pavillon E-4116 6875 LaSalle Blvd, Montreal (Verdun) QC H4H 1R3. Co-last authors (Véronique Fabre, Stéphanie Daumas).

## Abstract

The nucleus accumbens (NAc) is a key structure translating dopaminergic inputs into social avoidance following repeated social defeat. Within the NAc, dopamine (DA) release is modulated in part by cholinergic interneurons (CINs). CINs express the vesicular glutamate transporter type 3 (VGLUT3), which supports glutamate co-release and regulates vesicular acetylcholine (ACh) loading, thereby fine-tuning cholinergic signaling. A rare human VGLUT3-p.T8I missense variant is associated with compulsive behaviors toward food and drugs of abuse. We hypothesized that p.T8I disrupts NAc ACh dynamics and affects social avoidance following social defeat. Male and female mice expressing the VGLUT3-p.T8I variant were exposed to a 10-day chronic social defeat stress (CSDS) paradigm. Social avoidance toward the aggressor strain and a C57BL/6 (B6) conspecific was measured using the social interaction and 3-chamber tests, and anxiety-like behavior was evaluated in the elevated zero maze. Fiber photometry was used to monitor ACh and DA dynamics in the NAc medial shell (NAcMsh) during CSDS and social interaction. After CSDS, VGLUT3^T8I/T8I^ males showed aggressor strain-specific social avoidance and increased anxiety-like behavior compared to WT mice. Mutants showed blunted CSDS-induced ACh increase during attacks but enhanced ACh release upon re-exposure to the CD1 aggressor strain correlating with avoidance severity. DA dynamics were minimally affected. Females, regardless of genotype, displayed a male-mutant-like phenotype. In males, the VGLUT3-p.T8I variant alters NAcMsh ACh signaling and promotes selective social avoidance toward aggressors and elevated anxiety after CSDS, highlighting a potential cholinergic mechanism underlying stress-induced social avoidance with broader relevance to psychiatric disorders.

## Introduction

Exposure to stress, particularly social stressors, increases the risk of developing psychiatric conditions such as generalized anxiety and major depressive disorder [1–3]. However, the emotional and behavioral consequences of stress are highly heterogeneous, reflecting the complex interplay between genetic factors and environmental experiences. Within this framework, resilience has been conceptualized to be an active and adaptive process through which individuals maintain or regain functional behavior despite adversity [1, 4–6]. Importantly, resilience does not simply reflect the absence of stress-related symptoms, but rather the capacity to proactively adapt behaviorally to changing environments even in their presence [6–9].

This behavioral adaptation is thought to emerge from decision-making processes orchestrated by the striatum, where dopamine (DA) and acetylcholine (ACh) signals interact in a complex and dynamic manner [10–16]. DA transmission is widely implicated in motivation, reward prediction, and approach behavior, and ACh, released by cholinergic interneurons (CINs), supports reward-based decision-making, associative learning and behavioral flexibility in response to environmental changes [12, 17, 18]. Within the nucleus accumbens (NAc), DA release and the activity of dopaminergic terminals arising from the ventral tegmental area (VTA) encode social stress-related experiences and are causally involved in stress susceptibility versus resilience [8, 19–22]. However, the role of ACh signaling within the NAc in shaping stress responses and resilience remains incompletely understood. Silencing neurotransmitter release from CINs promotes depressive-like behaviors, while manipulation of their activity modulates stress susceptibility [23, 24]. Yet, these approaches do not account for the ability of CINs to co-release ACh and glutamate. This dual neurotransmitter phenotype arises from the expression of both the vesicular acetylcholine transporter (VAChT) and the vesicular glutamate transporter 3 (VGLUT3) [25–28]. VGLUT3 not only allows glutamate release but also enhances ACh vesicular loading through vesicular synergy, thereby influencing the magnitude of cholinergic signaling [25]. Recently, we identified in a population of severe substance addiction, a rare allelic variant (p.T8I) in the gene encoding VGLUT3 (*Slc17a8*) that selectively reduces ACh release from CINs without altering glutamate release [16, 29]. Mice expressing the p.T8I mutation (VGLUT3^T8I/T8I^) exhibit diminished DA levels in the dorsomedial striatum and maladaptive, habit-like behaviors as well as eating and substance use disorders [16]. These findings suggest that the p.T8I mutation affects striatal mechanisms underlying behavioral rigidity. Building on this, we used the VGLUT3^T8I/T8I^ mouse line to test whether VGLUT3-dependant ACh transmission also contributes to behavioral adaptation under social stress. Male and female VGLUT3^T8I/T8I^ mice were exposed to chronic social defeat stress (CSDS), a well-established model of stress-related psychiatric disorders, and were assessed for social and anxiety-like behaviors [30–32]. We then used fiber photometry to test whether these behavioral outcomes were associated with changes in NAc ACh and DA dynamics during CSDS and the social interaction test.

## Material and Methods

### Animals

All procedures complied with European directive 2010/63/EU and French regulations (authorization #31718-2021051822188705; ethics committee Darwin CEEA #5). Homozygous VGLUT3^T8I/T8I^ mice and WT littermates (VGLUT3^+/+^) were bred at Sorbonne University and used for all comparisons. Male and female mice (8-16 weeks) were housed at 21°C ± 2°C, 40% humidity, under a 12h light/dark cycle (07:30-19:30), with ad libitum access to food and water. Adult male CD1 mice (12-20 weeks, 30-35 g; Janvier Laboratories, France) served as aggressors for CSDS. Juvenile C57BL/6 mice (6 weeks, sex-matched) were used as social cues in the 3-chamber test.

### Modified chronic social defeat stress

CSDS was adapted from established protocols [30, 31, 33]. WT and VGLUT3^T8I/T8I^ mice were exposed to a novel male CD1 aggressor each day for 10 consecutive days. Each session consisted of a 5-min direct physical interaction followed by 20 min of protected sensory contact through a perforated divider. Female mice had urine from an unfamiliar male CD1 mouse applied around the genitals immediately before the physical interaction [32, 34].

### Behavioral Testing

Behavioral tests were conducted before and after CSDS. The social interaction (SI) test was performed in an open arena (42x42x26 cm, 10 lux) with two 150-s trials: one with an empty enclosure and one with an unfamiliar CD1 mouse. The three-chamber test (3C; 40x20x22 cm, 15 lux, 10 min) assessed social preference toward a juvenile sex-matched conspecific confined in a wire cage versus an empty cage. Anxiety-like behavior was assessed in the elevated zero maze (EZM; annular platform, 7-cm width, 57-cm outer diameter, elevated 90 cm, 30 lux) over 10 min. All sessions were video-recorded and analyzed with EthoVision 14.0 (Noldus, Netherlands).

### Stereotaxic Surgeries

Mice were anesthetized with ketamine/xylazine (90/10 mg/kg i.p.) and received local lidocaine (5 mg/kg) and postoperative meloxicam (1 mg/kg). Coordinates were adapted from the mouse brain atlas [35]. For fiber photometry, AAV2/9-hsyn-DA3h and/or AAV2/9-hsyn-ACh3.8 (each mixed 1:1 with AAV5/2-hSyn1-mCherry; Brain VTA) were infused (0.5 µL per side) into the nucleus accumbens (NAc) medial shell (AP +1.4 mm, ML ± 1.6 mm, DV -4.6 mm, 10° angle) [36]. Optical fibers (200 µm, NA 0.37; Neurophotometrics, USA) were implanted above the injection sites and secured with charcoal-mixed dental cement. Mice recovered for 3 weeks before experiments.

### In Vivo Fiber Photometry

DA and ACh signals were recorded using a dual-channel fiber photometry system (FP3002; Neurophotometrics) with 470 nm and 560 nm excitation lights (50 µW each; 75 Hz), synchronized with video via Bonsai 2.4. Recordings were performed during the SI test before and after CSDS and on CSDS days 1 and 10. Raw signals were preprocessed in Python. After removal of the first 5 s of recording, the sensor (470 nm) and control (560 nm) signals were detrended using double-exponential fits. The detrended control signal was linearly regressed onto the detrended sensor signal to calculate ΔF/F. Z-scores were computed from ΔF/F and aligned to behavioral events to generate peri-event time histograms (PETHs) from which areas under the curve were and max z-score values were extracted for statistical analysis.

### Immunohistochemistry

At the end of fiber photometry experiments, mice were perfused with 0.9% NaCL followed by ice-cold 4% paraformaldehyde (PFA) under deep anesthesia (Euthasol, 400 mg/kg i.p.). Brains were sectioned at 35 µm on a vibratome (Leica), then incubated with anti-GFP (chicken, 1:2000; Aves Labs #1020) and anti-RFP (mouse, 1:1000; ChromoTek #5F8) primary antibodies, followed by appropriate fluorescent secondary antibodies. Sections were mounted with Fluoromount-G. Mice lacking viral expression or with incorrect targeting of the NAc medial shell were excluded from analyses.

### Statistical Analysis

Data were analyzed in Prism 10.3 (GraphPad) and Python. Normality was assessed with the Shapiro-Wilk test. Group comparisons were performed using unpaired t-tests or two-way (repeated-measures) ANOVA, with Sidak’s post hoc tests or uncorrected Fisher’s LSD when appropriate. Non-parametric comparisons were performed using Wilcoxon signed-rank tests or other rank-based tests as appropriate. Missing data resulting due to recording failures were handled using mixed-effects models. Logistic regression was used to estimate predicted probabilities and assess classification performance by ROC curve analysis. PCA was performed on z-scored behavioral variables. Associations between behavioral and photometric data were assessed using Spearman’s rank correlations. Statistical significance was set at p < 0.05. Additional details are provided in the Supplementary Methods.

## Results

### The p.T8I variant promotes threat-associated social discrimination and anxiety-like behavior following CSDS in male mice

The VGLUT3-p.T8I variant was identified in cohorts with severe substance use disorders and eating disorders, both frequently comorbid with stress- and mood-related psychiatric disorders [16, 29, 37, 38]. We therefore investigated whether VGLUT3-p.T8I influences social and anxiety-like behaviors following CSDS. Male and female WT and VGLUT3^T8I/T8I^ mice were evaluated using the social interaction (SI), 3-chamber (3C) and elevated zero maze (EZM) tests before and after CSDS (Figure 1A and Supplementary Figure 1A; Supplementary Table 1).

**Figure 1.**
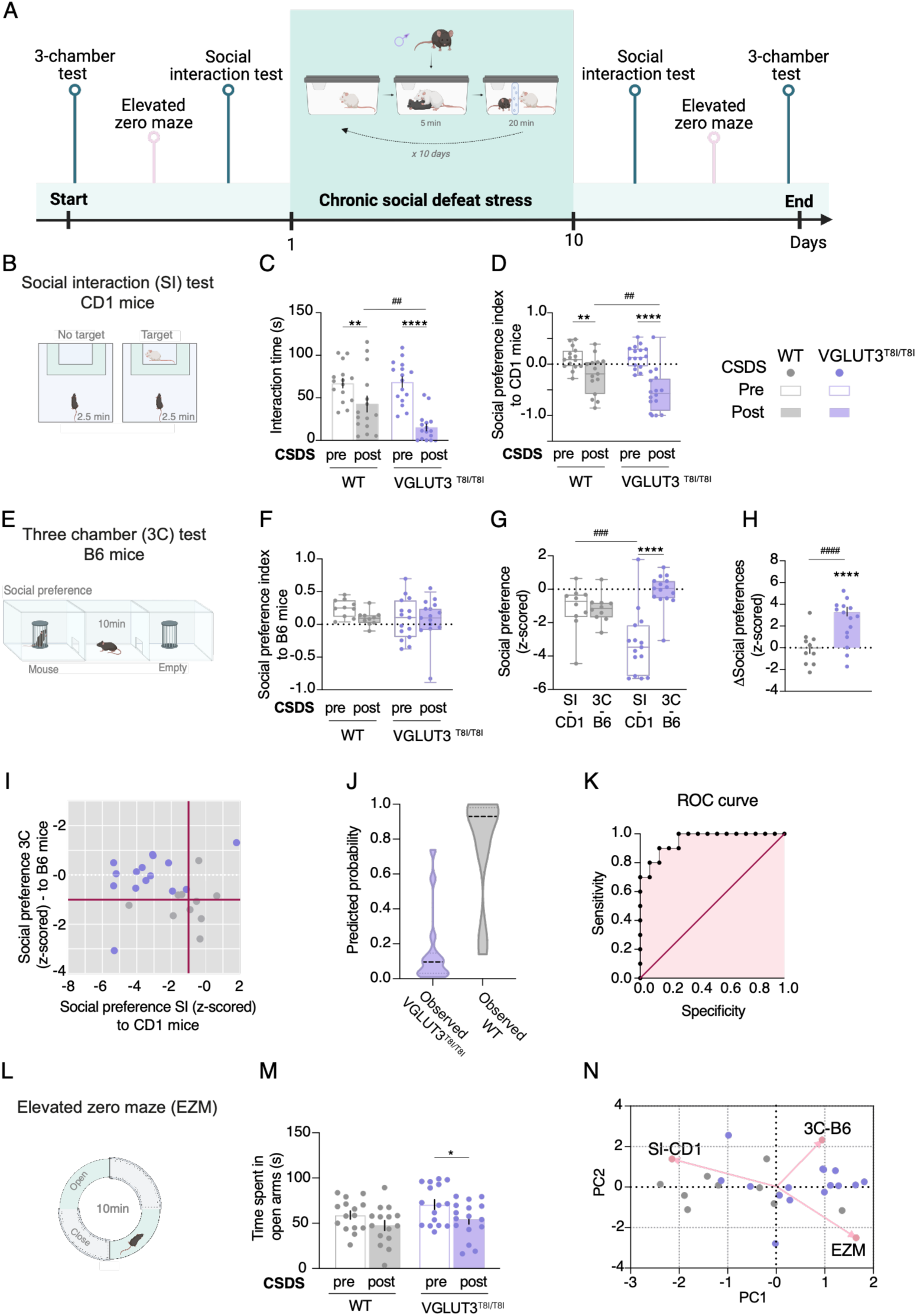
VGLUT3-p.T8I enhances threat-associated social discrimination after CSDS in male mice. **(A)** Experimental timeline. **(B)** Social interaction (SI) test schematic. **(C)** CSDS more markedly reduced SI interaction time in VGLUT3^T8I/T8I^ than WT mice. **(D)** Social preference (SP) indices toward the aggressor strain (CD1) significantly decreased in mutants post-CSDS relative to pre-CSDS in the SI test. **(E)** Three-chamber (3C) test schematic. **(F)** No significant CSDS or genotype effect on SP indices for a C57BL/6N conspecific (B6) in the 3C test. **(G)** Z-scored SP values for the CD1 and B6 mice revealed mutant-selective CD1 avoidance. **(H)** Mutants showed significantly greater delta changes in SP toward the CD1 aggressor strain than toward a B6 conspecific. **(I)** Scatter plot of z-scored SP values for the CD1 and B6 mice revealed distinct genotype-specific behavioral profiles. **(J)** Logistic regression using B6 vs CD1 SP robustly classified genotypes. **(K)** ROC curve confirmed model discriminative power (AUC=0.9533, p=0.0002). **(L)** Elevated zero maze (EZM) schematic. **(M)** VGLUT3^T8I/T8I^ mice spent significantly less time in open arms post-CSDS. **(N)** Principal component analysis (PCA) scores plot with PC1/PC2 loadings revealed distinct genotype-dependent behavioral clustering. WT mice, n=10-15 (grey bars); mutant mice, n=15-16 (purple bars); *p<0.05, **p<0.01, ****p<0.0001(CSDS effect), ^##^p<0.01, ^###^p<0.001, ^####^p<0.0001 (genotype effect; Sidak’s post-hoc, two-way RM ANOVA; one sample t test; unpaired t test).

Following CSDS, both WT and VGLUT3^T8I/T8I^ males showed reduced interaction time during the SI test compared to pre-CSDS, a reduction more pronounced in mutants (Figure 1B-C). Social preference (SP) indices, reflecting preference for the CD1 aggressor over the empty box, were also significantly decreased after CSDS in both genotypes, with a greater reduction in VGLUT3^T8I/T8I^ mice (Figure 1D). These results indicate that VGLUT3^T8I/T8I^ males display exacerbated social avoidance toward CD1 mice following CSDS. In the classical CSDS paradigm, mice are classified as susceptible or resilient based on whether they maintain post-CSDS preference for the CD1 mouse. Applying this criterion, 10/15 (66.67%) WT mice were classified as susceptible compared to 15/16 (93.75%) of VGLUT3^T8I/T8I^ mice.

To assess the specificity of social avoidance, we evaluated SP toward a B6 conspecific in the 3C test (Figure 1E-F). Neither CSDS nor genotype affected SP indices toward B6 mice, indicating that stress-induced social avoidance did not generalize to B6 conspecifics. To compare cue-specific changes across tests, SP indices were z-scored (Supplementary Methods) [39]. WT mice showed similar z-scored SP toward CD1 and B6 mice, whereas VGLUT3^T8I/T8I^ mice exhibited a pronounced reduction in SP toward the CD1 strain, with no change toward the B6 conspecific (Figure 1G). To further quantify the specificity of CSDS-induced changes, we computed the difference between z-scored SP indices obtained in the SI and 3C tests. Delta z-scored SP indices were near zero in WT mice, reflecting comparable responses across both social contexts (Figure 1H). In contrast, VGLUT3^T8I/T8I^ mice showed delta z-scored SP indices with values around 3, confirming selective attenuation of SP toward the CD1 strain. We then used scatter plots of z-scored SP indices to visualize these genotype-specific behavioral profiles. In Figure 1I, red threshold lines indicate significant deviations from baseline. Relative to pre-CSDS, most WT mice showed their strongest shift along the B6-related dimension, whereas VGLUT3^T8I/T8I^ mice deviated selectively toward the threat-associated CD1 cue with no change toward the B6 conspecific. A multiple logistic regression model based on z-scored SI and 3C SP indices classified genotype with 80% accuracy in WT mice and 86.7% in VGLUT3^T8I/T8I^ mice, with an area under the curve (AUC) of 0.9533 (Figure 1J-K). Higher SI scores (SP toward the CD1 strain) increased the odds of being WT, whereas higher 3C scores (SP toward the B6 conspecific) decreased them (Supplementary Table 1). Together, these results identify threat-associated social discrimination as a robust behavioral signature of the VGLUT3-p.T8I genotype.

Beyond social avoidance, VGLUT3^T8I/T8I^ males displayed heightened anxiety-like behavior post-CSDS, with reduced open-arm exploration in the EZM relative to WT mice (Figure 1L-M). This modest reduction likely reflected anxiety rather than motor impairment, since locomotion was unchanged in the SI and 3C tests (Supplementary Figure 1). Dark-light performance and grooming behavior in the splash test were also unaffected by CSDS or genotype, arguing against a broader anxiety-like or self-care deficit (Supplementary Figure 1). To integrate the behavioral dimensions assessed across tests, we performed a principal component analysis (PCA) on SI, 3C, and EZM data (Figure 1N). WT and mutant mice segregated primarily along the first principal component (PC1), with PC1 and PC2 together explaining 79.33 % of total variance. PC1 was positively associated with B6 preference and anxiety-like behavior and negatively associated with CD1 preference (Supplementary Table 1). Overall, VGLUT3^T8I/T8I^ mice showed a genotype-specific combination of heightened anxiety and selective social avoidance toward the CD1 strain, without generalization to the B6 conspecific.

The same behavioral battery was conducted in female mice (Supplementary Figure 2). Regardless of genotype, females displayed a male-mutant-like phenotype after CSDS, with selective avoidance of the CD1 strain and preserved SP toward the B6 conspecific. This indicates that sex strongly modulates the impact of the VGLUT3-p.T8I variant on stress-induced social behavior. In WT mice, CSDS also elicited sex-specific effects (Supplementary Figure 3). Indeed, WT females displayed a stronger anxiety-like phenotype and a larger decrease in SP toward the CD1 strain than WT males. In contrast, WT males showed only social avoidance and no increase in anxiety-like behavior. Overall, sex differences in this model were most evident in anxiety-like behavior and threat-associated social avoidance, rather than in general sociability.

### VGLUT3-p.T8I dysregulates CSDS-induced NAcMsh ACh dynamics in male mice

DA signaling in the NAc has been proposed to encode the valence of social experiences, and some studies suggest that ACh release from CINs drives DA release [8, 12, 16, 20, 27, 40]. Accordingly, we recorded ACh and/or DA signals in the NAc during social experiences in male WT mice and VGLUT3^T8I/T8I^ mice using fiber photometry. The NAc medial shell (NAcMsh) was targeted as it receives dense dopaminergic inputs from the VTA and plays a preferential role in aversion processing [41–43].

AAVs encoding the GACh3.8 and/othe GRAB-DA3h, along with a non-sensor mCherry control, were infused into the NAcMsh and optic fibers were implanted above the injection site (Figure 2A; Supplementary Figure 4A) [36].

**Figure 2.**
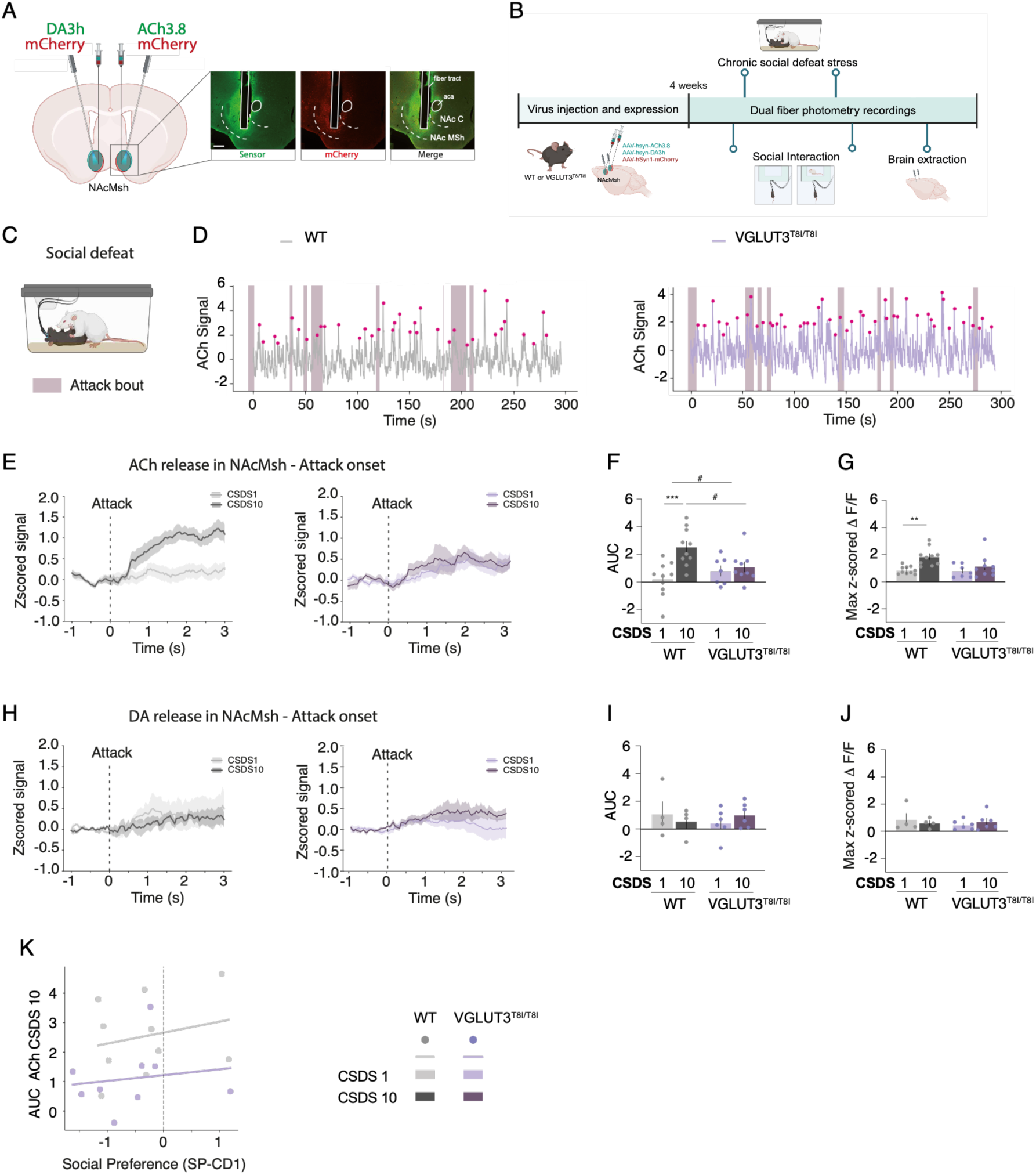
VGLUT3-p.T8I blunts chronic stress-induced ACh potentiation in the NAcMsh in male mice. **(A)** Schematic of surgical strategy for recording GRAB-DA3h and/or GACh3.8 sensors and mCherry as control in the NAcMsh. Representative fluorescence images depict sensor expression (green), mCherry (red), and merged signals with optic fiber placement (scale bar, 200 μm). Aca: anterior commissure; NAc: nucleus accumbens; NAc C: nucleus accumbens core ; NAcMSh: nucleus accumbens medial shell. **(B)** Experimental timeline for fiber photometry recordings. **(C)** Social defeat recording schematic. **(D)** Representative z-scored ΔF/F of ACh signal during CSDS in WT (grey) and VGLUT3^T8I/T8I^ mice (purple); shaded regions indicates attack bouts. **(E)** Peri-event traces of ACh signals aligned to attack onset by the CD1 mouse on CSDS day 1 and day 10 in WT (left) and VGLUT3^T8I/T8I^ (right) mice. **(F)** AUC of ACh signals significantly enhanced on CSDS day 10 relative to day 1 in WT mice but not in VGLUT3^T8I/T8I^ mice. **(G)** Peak z-scored ΔF/F of ACh signals higher on CSDS day 10 in WT mice but not in VGLUT3^T8I/T8I^ mice. **(H)** Peri-event traces of DA signals aligned to attack onset by the CD1 mouse on CSDS day 1 and day 10 in WT (left) and VGLUT3^T8I/T8I^ (right) mice. **(I)** AUC of DA signals unaffected across CSDS days in both genotypes. **(J)** Peak z-scored ΔF/F of DA signals showed no significant day or genotype effect. **(K)** AUC ACh signals during the 3 s following attack onset [0-3 s] on CSDS day 10 did not correlate with subsequent social preference (SP) scores toward the CD1 aggressor strain evaluated in the SI test (WT: ρ=0.12; VGLUT3^T8I/T8I^: ρ=0.32; Spearman’s rank correlation). *p<0.05, **p<0.01 (CSDS effect; mixed-effects analysis with Sidak post hoc comparisons).

ACh and DA signals were recorded on the first (CSDS1) and the last (CSDS10) days of the CSDS and during the SI tests pre- and post-CSDS (Figure 2B). Signals were aligned to attack onset by the CD1 mouse on CSDS1 and CSDS10 (Figure 2C-D). WT mice showed stress-induced enhanced ACh reponses between CSDS1 to CSDS10, as evidenced by increased mean AUC and peak z-scored ACh signals at attack onset on CSDS10 (Figure 2E-G). This augmented ACh efflux was absent in VGLUT3^T8I/T8I^ mice, suggesting that the VGLUT-p.T8I variant blunts the ACh response to repeated social defeat. DA signals were unchanged across CSDS sessions in both genotypes (Figure 2H-J), indicating differential regulation of ACh and DA dynamics in the NAcMsh during chronic stress. Cross-correlation analyses further showed unaffected ACh/DA temporal coupling at attack onset in both genotypes, with similar peak correlations near zero lag on CSDS1 and CSDS10 (Supplementary Figure 4B-C). To assess whether ACh dynamics during social defeat predict subsequent social behavior toward the CD1 aggressor strain, we correlated AUC of ACh signals during CSDS10 attacks with z-scored SP indices in the post-CSDS SI test (Figure 2K). ACh signals did not correlate with post-CSDS SP indices in either genotype, suggesting that stress-induced increase in ACh during CSDS is dissociable from later social avoidance toward CD1 mice (Supplementary Table 1).

ACh and DA signals in the NAcMsh were analyzed during the SI test by aligning activity to nose-to-nose interactions with the CD1 mouse before and after CSDS (Figure 3). In WT mice, neither ACh nor DA efflux changed significantly between pre- and post-CSDS conditions (Figure 3A-F). In contrast, VGLUT3^T8I/T8I^ mice displayed a significant increase in ACh signals during post-CSDS social interaction with the CD1 strain, whereas DA signals were not significantly altered (Figure 3A-F). Cross-correlation analyses further showed preserved ACh/DA coupling in both genotypes before and after CSDS (Supplementary Figure 4D-E).

**Figure 3.**
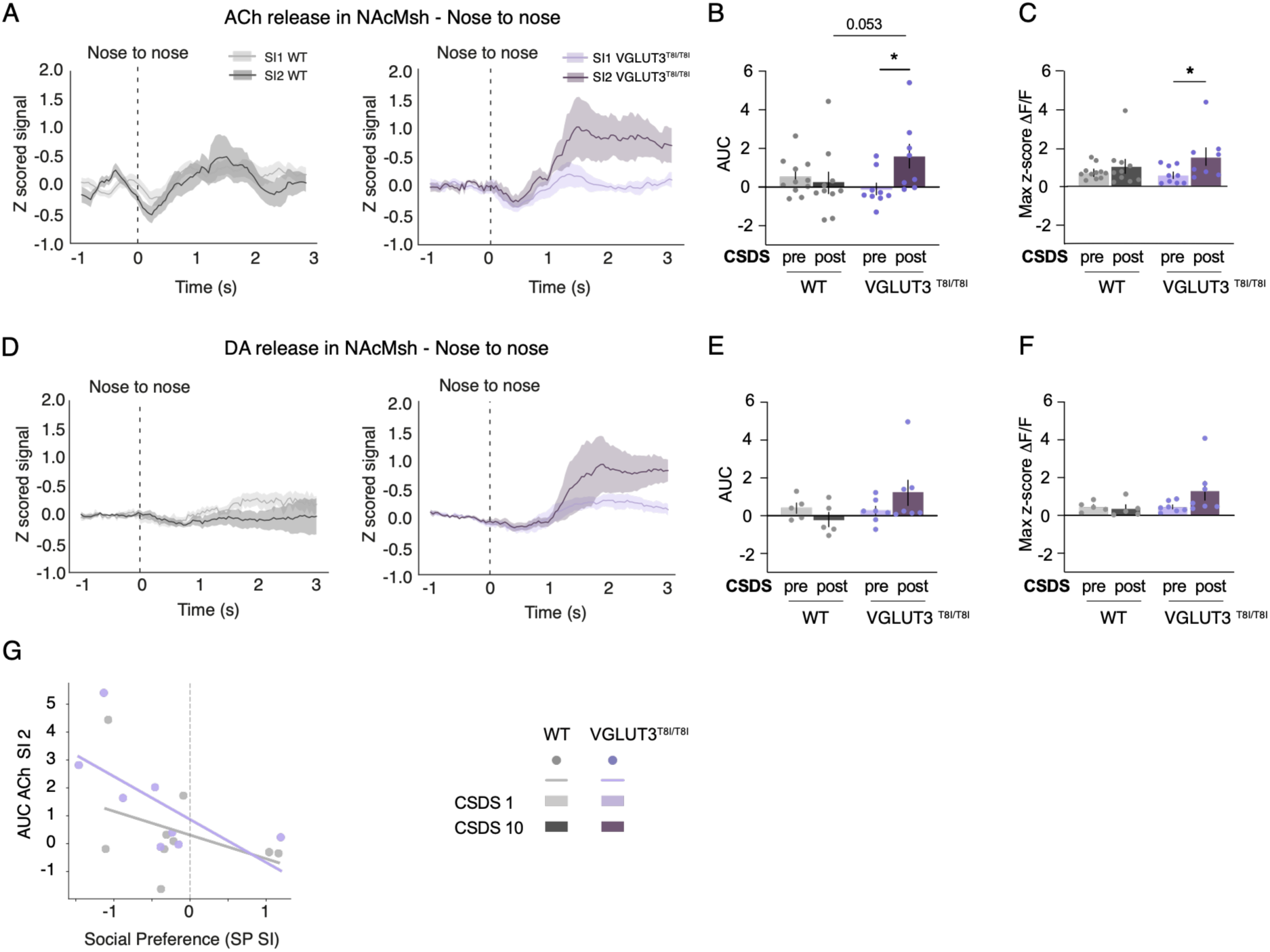
The VGLUT3-p.T8I variant induces exaggerated NAcMsh ACh signals in male mice during post-stress social interaction, correlating with social avoidance. **(A)** Peri-event traces of ACh signals aligned to nose-to-nose interactions with the CD1 mouse pre- and post-CSDS during the social interaction (SI) test in WT (left) and VGLUT3^T8I/T8I^ (right) mice. **(B)** AUC of ACh signals significantly enhanced post-relative to pre-CSDS in VGLUT3^T8I/T8I^ mice but no in WT mice. **(C)** Mean peak z-scored ΔF/F of ACh signals showed the same pattern. **(D)** Peri-event traces of DA signals aligned to nose-to-nose interactions with the CD1 mouse pre-and post-CSDS during the SI test in WT (left) and VGLUT3^T8I/T8I^ (right) mice. **(E)** AUC of DA signals unaffected by CSDS in both genotypes. **(F)** Peak z-scored ΔF/F of DA signals showed no significant CSDS or genotype effect. **(G)** Post-CSDS SI AUC ACh signals during the 3 s following interaction onset [0-3 s] negatively correlated with social preference (SP) in mutants (ρ=−0.79, P=0.021; Spearman’s rank correlation). *p<0.05 (CSDS effect; mixed-effects analysis with Sidak post hoc comparisons; Spearman’s rank correlation).

However, VGLUT3^T8I/T8I^ mice showed a negative peak lag, indicating that DA fluctuations preceded ACh changes (Supplementary Figure 4D-E). To determine whether ACh dynamics in the SI test were associated with social avoidance, we correlated AUC of ACh signals and z-scored SP indices during the post-CSDS SI test (Figure 3G). In WT mice, ACh signals were not significantly correlated with SP indices (Supplementary Table 1). In VGLUT3^T8I/T8I^ mice, ACh signals during social interaction with the CD1 strain were negatively correlated with SP indices (Supplementary Table 1).

Together, these findings reveal a differential dynamic of ACh in the NAcMsh of WT mice and VGLUT3^T8I/T8I^ mice during social defeat and post-stress social interaction. WT mice show CSDS-induced augmentation of ACh signaling at attack onset on the last day of CSDS. In contrast, VGLUT3^T8I/T8I^ mice fail to develop this adaptation and instead show increased ACh signals during post-CSDS exposure to the CD1 strain. Notably, the magnitude of this response correlates with social avoidance. These results suggest that VGLUT3-dependent cholinergic signaling in the NAcMsh contributes to the encoding of threat-associated social cues following CSDS.

## Discussion

Understanding how neurotransmitter dynamics shape adaptive versus maladaptive responses to stress remains a central question with important implications for stress-related psychiatric disorders. While DA signaling in the NAc has been linked to stress susceptibility [8, 19, 44], the contribution of ACh released by CINs remains poorly understood [23, 24]. CINs co-express VGLUT3, which enables glutamate release and increased vesicular ACh uptake through vesicular synergy [25, 26]. The human VGLUT3-p.T8I variant does not affect glutamate release but selectively impairs vesicular synergy and reduces ACh release [16]. This mutation thereby provides a unique opportunity to investigate striatal cholinergic contributions to stress-induced behavioral adaptations. Using the CSDS paradigm, we show that male mice carrying the VGLUT3-p.T8I variant display strain-specific social avoidance toward the threat-associated CD1 strain. In addition, the p.T8I mutation increases anxiety and alters NAcMsh ACh dynamics during aggressor attacks and subsequent CD1 exposure in the SI test. Regardless of their genotype females also showed strain-selective avoidance and heightened anxiety. Together, these findings reveal that the VGLUT3-p.T8I variant produces a male-specific strain-selective avoidance phenotype associated with specific NAcMsh ACh dynamics during social threat exposure. Our study thus highlights a previously underappreciated role for cholinergic transmission in shaping stress-induced behavioral adaptations.

In VGLUT3^T8I/T8I^ males, CSDS induced pronounced social avoidance toward the threat-associated CD1 strain, together with increased anxiety-like behavior. Although the conventional SI ratio would classify this phenotype as increased stress susceptibility, responses toward safe-associated conspecifics indicate preserved threat-safety discrimination rather than generalized social withdrawal [45]. Several studies suggest that avoidance directed specifically toward the aggressor strain may reflect adaptive encoding of threat rather than maladaptive social impairment [45, 46]. By contrast, WT males exhibited poorer discrimination between threat- and safe-associated strains, consistent with a more generalized social avoidance phenotype [39, 45, 46]. This enhanced strain-specific discrimination fits with a broader phenotype associated with the VGLUT3-p.T8I variant. In a previous study, this variant shifted the balance away from goal-directed control toward habitual responding and facilitated reinstatement after extinction, consistent with strengthened stimulus-response associations [16]. These behaviors are generally considered as maladaptive because they reduce behavioral flexibility. However, they also reflect an enhanced capacity to form persistent cue-specific associations. Together with the present findings, they suggest that the VGLUT3-p.T8I variant biases learning toward more rigid cue-outcome encoding across aversive and appetitive domains. In the context of social defeat, this may promote more precise discrimination between threatening and safe social cues and limit the generalization of avoidance to neutral conspecifics.

Male odor pre-exposure in the adapted CSDS protocol enables the assessment of stress-induced behavioral outcomes in females and direct comparison with males [32, 34]. Consistent with previous female-adapted CSDS studies [32, 34], defeated WT females showed selective social avoidance toward the CD1 aggressor strain together with increased anxiety-like behavior. Social avoidance remained restricted to the treat-associated CD1 strain and did not generalize to safe-associated B6 conspecifics, indicating preserved threat-safety discrimination. This behavioral profile contrasts with that of WT males, which exhibited a more generalized threat response, and is consistent with previous reports of sex differences in aversive learning and fear generalization [47]. In contrast to males, defeated female mutants did not differ from controls in social avoidance, anxiety-like behavior, or threat discrimination, suggesting that the VGLUT3-p.T8I variant does not alter stress-induced behavioral responses in females. These findings indicate a sex-specific effect of the variant, suggesting differential regulation of VGLUT3 function across sexes. Whether this sexual dimorphism also extends to reward-related behavioral alterations previously described in male mutants remains to be determined [16].

To investigate the neural dynamics underlying the enhanced, strain-specific social avoidance displayed by mutant males, we monitored ACh and DA dynamics in the NAcMsh during repeated aggressor attacks across CSDS and subsequent exposure to a CD1 mouse in the SI test. In WT males, attack-induced ACh release increased from day 1 to day 10 of CSDS, indicating a progressive cholinergic potentiation across repeated social defeat. This increase is consistent with the growing view that CSDS involves associative aversive learning, in which repeated encounters strengthen the association between the aggressor cue and the negative consequences of attack [45, 46]. Across reward and aversive learning, ACh is thought to regulate associative representations rather than directly encode reinforcement value [48–50]. Furthermore, cue-evoked ACh signals in the ventral striatum progressively emerge as learning develops [51]. Thus, the ACh increase in WT mice likely reflects reinforced updating of the CD1 threat representation. In contrast, VGLUT3^T8I/T8I^ males failed to exhibit this stress-dependent ACh potentiation. Given that VGLUT3-p.T8I variant selectively impairs vesicular ACh loading and release [16], this blunted scaling suggests a reduced capacity of cholinergic signaling to adapt to repeated social defeat, potentially limiting the amplification of aversive processing [48, 52] and activity-dependent cholinergic receptor desensitization. Importantly, ACh dynamics on day 10, measured as AUC during the attack, did not correlate with subsequent social avoidance in the SI test, indicating that cholinergic responses during defeat alone do not determine the later behavioral phenotype. A markedly different pattern emerged during the re-exposure to the aggressor strain. Following CSDS, only VGLUT3^T8I/T8I^ males showed a significant increase in ACh AUC during approach to the social threat after CSDS, which negatively correlated with the social preference index. This indicates that greater ACh release was associated with stronger social avoidance in mutants, in line with studies linking striatal ACh with aversive learning [48, 52], threat cue processing [23, 24, 52, 53] and associative salience during learning [49, 53, 54]. Therefore, increased ACh release in the NAcMsh of VGLUT3^T8I/T8I^ males may reflect enhanced aversive salience of CD1 mice during re-exposure which may promote selective avoidance of the aggressor strain while sparing interactions with neutral conspecifics [53, 55, 56]. Together, these findings suggest that the VGLUT3-p.T8I variant alters the temporal dynamics rather than the overall engagement of cholinergic signaling during stress. By preventing the progressive hyper recruitment of NAcMsh cholinergic activity during CSDS, the variant may preserve the capacity for ACh response during later threat re-exposure. Such a mechanism would favor precise encoding of cue-specific threat salience [48, 56] and adaptive avoidance of the aggressor strain [45, 55] while limiting the generalization of defensive responses to safe social stimuli [50, 57].

In contrast to ACh, DA dynamics in the NAcMsh remained stable across CSDS and during the SI test in both genotypes, consistent with the idea that dopaminergic responses are topographically regulated within the NAc [41, 58, 59] and may differ from reports focusing on other NAc subregions [8, 20]. Although overall DA responses were unaffected by genotype, cross-correlation analyses suggest that VGLUT3-p.T8I alters DA-ACh timing: coupling was modest and unbiased during CSDS10, but VGLUT3^T8I/T8I^ males showed a more consistent DA-before-ACh relationship during the SI test. Thus, the variant may not change dopaminergic recruitment per se, but instead reshape DA-ACh coordination during social behavior, potentially biasing cue processing toward a more stereotyped pattern [13, 40, 60–62].

The VGLUT3-p.T8I variant, which reduces vesicular synergy and ACh exocytosis in striatal CINs, was first identified in patients with severe addiction [16, 29]. Here, it promoted cue-specific rather than generalized social avoidance after CSDS in male mice. Mutant mice preserved discrimination between threat- and safe-associated conspecifics, suggesting enhanced encoding of socially relevant threat cues. This phenotype was accompanied by altered ACh dynamics in the NAcMsh, with reduced stress-induced ACh potentiation during repeated social defeat and enhanced ACh responses during subsequent re-exposure to the aggressor strain. Interestingly, the magnitude of this ACh responses predicted social avoidance. These findings suggest that VGLUT3-dependent co-transmission shapes accumbal cholinergic signaling during social stress, influencing the balance between threat discrimination and threat generalization. Altered NACMsh cholinergic dynamics may therefore contribute to context-appropriate social threat encoding. Because the VGLUT3-p.T8I variant appears to limit maladaptive social threat generalization, pro-cholinergic strategies may improve threat discrimination in stress-related psychiatric disorders.

## Acknowledgments and Disclosures

We thank the Institut de Biologie Paris-Seine (IBPS) Animal facility for breeding and care (CEFI), the NeuroSU Phenotypic core facility, especially Jean Vincent; Jean-François Gilles and France Lam from the IBPS Photon microscopy facility; Aquineuro and especially Sebastien Delcasso and Rémi Proville for their valuable scientific insights and support for fiber photometry pipeline analysis; and undergraduate students Khadija Laurent-Haond and Lucille Desvignes, for technical help.

Figures created with BioRender.com.

## Author contributions

S.D. acquired funding. M.P., V.F., and S.D. conceptualized the study. M.P. performed the behavioral and fiber photometry experiments and V.B. and V.F. acquired the microscopy images. M.P., V.F., and S.D. analyzed the data. M.P. drafted the manuscript. M.P., V.F., S.D., and S.E.M. reviewed and edited the manuscript. M.P., V.F., S.D, and S.E.M. prepared the figures.

## Conflict of interest

All authors have no relevant conflicts to declare.

Supplementary information is available at MP’s website

