## Supplementary Information for "The VGLUT3-p.T8I Mutation Alters Striatal Acetylcholine Dynamics in Response to Social Stressors and Drives Selective Social Avoidance in Male Mice"

**This PDF file includes:**

Supplementary Methods

Supplementary Figures 1-4

Supplementary Table 1

Supplementary References

#### SUPPLEMENTARY METHODS

##### Animals

All experiments were conducted in accordance with the European directive on the protection of animals used for scientific purposes (European Committee Council Directive 2010/63/EU) and complied with the regulations of the French Ministère de l'Agriculture et de la Forêt, Service Vétérinaire de la Santé et de la Protection Animale (authorization #31718-2021051822188705; ethics committee Darwin CEEA #5). WT and VGLUT3$^{T8I/T8I}$ mice were bred in the Sorbonne University rodent facility. Mice were housed under standard conditions at 21°C ±2°C, 40% humidity, and a 12h light/dark cycle (07:30-19:30), with food and water available ad libitum. Experiments were performed in male and female mice aged 8-16 weeks. Adult male CD1 mice (12-20 weeks-old, 30-35g; Janvier Laboratories, France) were used as aggressors for chronic social defeat stress (CSDS). For the 3-chamber test, juvenile C56BL/6 mice (6 weeks old, sex-matched) were used as social cues. The p.T8I allelic variant within the *Slc17a8* gene on a C57BL/6N background was used throughout the study (31). Homozygous VGLUT3$^{T8I/T8I}$ mice and WT littermate VGLUT3$^{+/+}$ mice were compared at the behavioral and biochemical levels.

##### Modified chronic social defeat stress

CD1 male aggressors were pre-screened for aggressive behavior according to published criteria (1). A modified CSDS protocol was used to permit testing of female mice (2,3). Each day, the experimental mouse was introduced into the home cage of a novel CD1 aggressor for a 5-min physical interaction. This was followed by a 20-min period of protected sensory contact, during which the two mice were separated by a perforated Plexiglas partition permitting visual, olfactory, tactile, and auditory contact. After each 25-min session, experimental mice were isolated in their home cages. The cycle was repeated for 10 consecutive days (days 1-10), with a new aggressor each day.

For female mice, the protocol was adapted as follows (4,5): approximately 50 µL of urine from a non-aggressive male CD1 mouse was applied around the genitals of each female experimental mouse immediately before the 5-min physical interaction. This facilitated attack initiation by the CD1 aggressor.

#### Behavioral Testing

##### *Social Interaction test*

Social approach or avoidance was assessed using a two-trial social interaction test. In the first 2.5-min trial, the experimental mouse explored a white Plexiglas open arena (42x42x26 cm, 10 lux) containing an empty transparent perforated Plexiglas box (10x7x15 cm; "No target" condition). In the second 2.5-min trial, an unfamiliar male CD1 mouse was confined in the box while the experimental mouse explored the arena ("Target" condition). Time spent in the interaction zone (8-cm radius around the box) was recorded using EthoVision 14.0 (Noldus, Netherlands).

##### *Three-Chamber test*

Social preference toward a C57BL/6N conspecific was assessed in an opaque Plexiglas three-chamber apparatus (40x20x22 cm, 15 lux), with three compartments interconnected by 5x5 cm openings. After a 10-min habituation phase in the central chamber with doors closed, the experimental mouse was allowed to freely explore all three chambers for 10 min. An unfamiliar sex-matched juvenile mouse (C57BL/6N, 6 weeks old) was confined in a small, weighted wire cage in one chamber, while the other chamber contained an empty identical wire cage. Time spent in close interaction with each cage was quantified using EthoVision 14.0.

Across both social tests, social preference indexes were calculated as: Ix = (time near mouse − time near empty cage) / (time near mouse + time near empty cage). To assess the effect of stress, preference indexes were z-scored relative to pre-CSDS group values: z = (individual data − mean[pre-CSDS group values]) / SD[pre-CSDS group values]. These z-scores allowed the position of each individual after stress to be interpreted relative to the pre-CSDS group distribution. Animals were excluded if they exhibited abnormal test behavior, failed predefined validity criteria for the three-chamber assay, or could not be analyzed because of technical failures or death before testing.

##### *Elevated Zero Maze*

Anxiety-like behavior was assessed in an elevated zero maze (EZM), consisting of a white polyvinyl chloride (PVC) annular platform (width 7 cm, outer diameter 57 cm) elevated 90 cm above the floor, divided into two opposing open sections and two opposing closed sections (wall height 17 cm, 30 lux in open areas). Mice were placed in a closed section facing the open area and allowed to freely explore for 10 min. Total time spent in open sections were tracked using EthoVision 14.0.

**Stereotaxic Surgeries**

All animals were anesthetized with an intraperitoneal mixture of ketamine/xylazine (90 and 10 mg/kg, respectively) diluted in 0.9% NaCl and placed in a stereotaxic frame (David Kopf Instruments). Local analgesia was provided by lidocaine (5 mg/kg, applied topically), and postoperative systemic analgesia by meloxicam (1 mg/kg s.c., administered at the end of surgery and the following day).

**Fiber Photometry: Viral Vectors and Fiber Implantation**

AAV2/9-hsyn-DA3h ($5.85\times10^{12}$ vg/mL; Brain-VTA, Wuhan, China) and AAV2/9-hsyn-ACh3.8 ($2.05\times10^{12}$ vg/mL; Brain-VTA, Wuhan, China) were each mixed at a 1:1 ratio with AAV5/2-hSyn1-mCherry ($3\times10^{12}$ vg/mL; Viral Vector Facility, University of Zurich, Switzerland). The two viral mixtures were injected separately in male mice, with the injection side alternated between animals to prevent lateralization bias. A volume of 0.5 µL per mixture was bilaterally infused into the NAc shell at the following coordinates relative to bregma: AP +1.4 mm, ML ±1.6 mm, DV -4.6 mm (10° angle), at a flow rate of 0.2 µL/min using a 10 µL Hamilton syringe (model 1701) fitted with a silica fiber-tip needle (100 µm; PHYMEP, Paris, France). The needle was left in place for 5 min after infusion before withdrawal. Optical fibers (200 µm core diameter, 1.25-mm ceramic ferrule, NA 0.37, 7.5 mm length; Neurophotometrics, San Diego, USA) were implanted 0.2 mm above the injection site (DV -4.4 mm) and secured with charcoal-mixed dental cement (SuperBond, PHYMEP, Paris, France) to minimize ambient light interference. Mice recovered for 3 weeks to allow for adequate viral expression. Correct targeting was confirmed post-hoc by immunohistochemistry; mice lacking detectable viral expression or with misplaced fibers were excluded.

**In Vivo Fiber Photometry: Data Acquisition**

Fluorescent signals from DA3h, ACh3.8 biosensors and mCherry control were recorded simultaneously using a multi-fiber photometry system (FP3002; Neurophotometrics, USA) coupled to a branching low-autofluorescence patch cord (Doris Lenses). Two LED excitation lights (470 nm for biosensor signal, 560 nm for control, 50 µW each) were bandpass filtered, reflected by a dichroic mirror, and focused through a x20 objective (NA 0.4) before transmission to the implanted fiber. Signals were acquired at 75 Hz and synchronized with video recording via Bonsai 2.4 (http://bonsai-rx.org). One week prior to recordings, mice were habituated to the fiber patch cord connection. Recordings were performed during the SI test before and after CSDS, and on CSDS days 1 and 10.

**In Vivo Fiber Photometry: Signal Preprocessing**

All photometry data were analyzed in Python. The first 5 s of each recording were discarded to remove acquisition artifacts. The 470 nm sensor signal and the 560 nm control signal were each fitted with double exponential functions using nonlinear least squares, and each signal was detrended by dividing by its fitted curve. The detrended 560 nm control signal was then regressed onto the detrended 470 nm signal by ordinary least squares to yield a 'fitted control'. The change in fluorescence was computed as: $\Delta F/F$ = (sensor signal −fitted control) / fitted control. Z-scores were calculated using $z = (\Delta F/F - \text{mean}[\Delta F/F]) / \text{SD}[\Delta F/F]$. The z-scored signals were time-locked to behavioral events to generate peri-event time histograms (PETHs). The area under the curve (AUC) and max z-score of each PETH were extracted for statistical comparisons. Cross-correlation analyses were performed on z-scored ACh and DA signals to quantify their temporal coupling and relative lag structure.

**Immunohistochemistry**

Mice were anesthetized by intraperitoneal injection of Euthasol (400 mg/kg) and perfused intracardially with 100 mL of 4 % paraformaldehyde (PFA) solution buffered in phosphate-buffered saline (PBS). Brains were then removed and kept at 4°C in a 4 % PFA solution for 24 h before being transferred to a PBS solution with azide added (0.03 %). Coronal sections (35 µm) were cut on a vibratome (VT-1000, Leica Microsystems, Rueil-Malmaison, France) and stored at 4°C. Brain slices were then rinsed and incubated in a 4 % normal horse serum (NHS) blocking buffer diluted in 1x PBS and 0.3 % Triton for one hour. Afterwards, the sections were incubated overnight at 4°C with primary antibodies against GFP (1/1000; chicken; AVES) and against DSred (1/1000; rabbit; Abcam) diluted in a solution of 1x PBS, 0.3 % Triton, and 1 % NHS. After PBS washes, sections were incubated for 1 h with secondary antibodies anti-chicken conjugated to GFP (1/2000; Alexa Fluor 488 nm) and anti-rabbit conjugated to DSred (1/2000; Alexa Fluor 594 nm) diluted in 1x PBS. Then, sections were mounted between a slide and a coverslip using Fluoromount (Sigma-Aldrich, F4680) and stored at 4°C.

**Image Acquisition**

Dual-fluorescence images were acquired using a Zeiss Axio Zoom V16 microscope. For illustration, images were exported in TIFF format, adjusted for contrast, cropped, and assembled using Photoshop CS2 (version 9.0; Adobe Systems, Mountain View, CA, USA). Anatomical landmarks and other annotations were added for presentation purposes only.

**Statistical Analysis**

Data are presented as mean ± SEM, except for box-and-whisker plots, which show the median, minimum-to-maximum whiskers. Data were analyzed using Prism 10.3 (GraphPad) and Python. Normality was assessed using the Shapiro-Wilk test. Parametric data were compared using unpaired Student's t-tests, one-sample t tests, or two-way (repeated-measures) ANOVA; non-parametric data were analyzed using Wilcoxon signed-rank tests or appropriate rank-based tests. Post hoc analyses were performed using Sidak's multiple-comparisons test or uncorrected Fisher's LSD, as appropriate. When data were missing (e.g., due to photometry recording failures), mixed-effects models replaced repeated-measures ANOVA. Spearman correlations assessed monotonic relationships between behavioral and photometric variables. Cross-correlation analyses quantified temporal coupling between z-scored DA and ACh signals. Principal component analysis (PCA) was performed in Prism 10.3 on z-scored behavioral data (EZM, 3C, SI) normalized to pre-CSDS group values: z = (data − mean[pre-CSDS group values]) / SD pre-CSDS group values]. The proportion of variance explained by each PC and the variable contributions were also computed. Statistical significance threshold was set at $p < 0.05$.

### SUPPLEMENTARY TABLE 1. Statistical results for figures.

| Figure | n | Statistical analyses | | Value | p-value | Post-hoc test |
|---|---|---|---|---|---|---|
| **Fig. 1C** | WT : n=15<br>VGLUT3$^{T8I/T8I}$: n=16 | Interaction Time | | | | |
| | | Two-way RM ANOVA | Interaction | F (1, 29) =7.125 | =0.0123 | Uncorrected Fisher's LSD |
| | | | Genotype | F (1, 29) =3.043 | =0.0917 | |
| | | | Stress | F (1, 29) =51.62 | <0.0001 | |
| **Fig.1D** | WT: n=15<br>VGLUT3$^{T8I/T8I}$ : n=16 | Social preference index (CD1) | | | | |
| | | Two-way RM ANOVA | Interaction | F(1,29) = 6.718 | =0.0148 | Uncorrected Fisher's LSD |
| | | | Genotype | F(1,29) = 2.452 | =0.1282 | |
| | | | Stress | F(1,29) = 55.88 | <0.0001 | |
| **Fig.1F** | WT : n=10<br>VGLUT3$^{T8I/T8I}$ : n=15 | Social preference index (B6) | | | | |
| | | Two-way RM ANOVA | Interaction | F(1,23) = 1.428 | =0.2442 | |
| | | | Genotype | F(1,23) = 1.227 | =0.2795 | |
| | | | Stress | F(1,23) = 2.737 | =0.1116 | |
| **Fig.1G** | WT : n=10<br>VGLUT3$^{T8I/T8I}$ : n=15 | Social preference (z-scored) | | | | |
| | | Two-way RM ANOVA | Interaction | F(1,23) = 22.54 | <0,0001 | Uncorrected Fisher's LSD |
| | | | Genotype | F(1,23) = 1.433 | =0.2435 | |
| | | | Test | F(1,23) = 21.18 | =0.0001 | |
| **Fig.1H** | WT : n=10<br>VGLUT3$^{T8I/T8I}$ : n=15 | Delta social preferences (z-scored) | | | | |
| | | Unpaired t test two-tailed (WT vs. VGLUT3$^{T8I/T8I}$) | | t = 4.748 et DF = 23 | <0.0001 | |
| | | One sample t test (WT, VGLUT3$^{T8I/T8I}$) | | t = 0.09986 et DF = 9<br>t = 7.145 et DF = 14 | =0.9226<br><0.0001 | |
| **Fig. 1J-K** | VGLUT3$^{+/+}$ : n=10 | Multiple logistic regression (dependant variable = genotype) | | | | |

| | | | | | | |
|---|---|---|---|---|---|---|
| | VGLUT3$^{T8I/T8I}$ : n=15 | Odds ratios estimate | Intercept<br>SI<br>3C<br>SI:3C | Estimate = 2.088<br>Estimate = 2.490<br>Estimate = 0.07430<br>Estimate = 0.7347 | | |
| | | Hosmer-Lemeshow test | | | =0.8152 | |
| | | Tjur's R squared | | =0.6135 | | |
| | | ROC Curve | | Area = 0.9533 | =0.0002 | |
| | | Negative predictive power (%) | | =86.67 | | |
| | | Positive predictive power (%) | | =80.00 | | |
| **Fig.1M** | VGLUT3$^{+/+}$ : n=15<br><br>VGLUT3$^{T8I/T8I}$ : n=16 | Time spent in open arms (EZM) | | | | |
| | | Two-way RM ANOVA | Interaction | F(1,29) = 0.277 | =0.6022 | Sidak's multiple comparisons test |
| | | | Genotype | F (1,29) = 2.677 | =0.1126 | |
| | | | Stress | F(1,29) = 8.862 | =0.0058 | |
| **Fig.1N** | VGLUT3$^{+/+}$ : n=10<br><br>VGLUT3$^{T8I/T8I}$ : n=15 | Principal Component Analysis (PCA) | | | | |
| | | Eigenvalue | PC1 | 1,413 | | |
| | | | PC2 | 0,9667 | | |
| | | | PC3 | 0,6201 | | |
| | | Proportion of variance | PC1 | 47.11% | | |
| | | | PC2 | 32.22% | | |
| | | | PC3 | 20.67% | | |
| | | Loading values PC1 | SI | -0.7187 | | |
| | | | 3C | 0.4702 | | |
| | | | EZM | 0.8219 | | |
| | | Loading values PC2 | SI | 0.5010 | | |
| | | | 3C | 0.8447 | | |
| | | | EZM | -0.0451 | | |

| | | | | | | |
|---|---|---|---|---|---|---|
| **Fig. 2F** | WT : n=10<br>VGLUT3$^{T8I/T8I}$ : n=7-9 | AUC ACh Attack | | | | |
| | | Mixed-effects model (REML) | Genotype | F(1,18) = 0.9333 | =0.3468 | Uncorrected Fisher’s LSD |
| | | | Stress | F(1,14) = 11.83 | =0.0040 | |
| | | | Interaction | F(1,14) = 7.288 | =0.0173 | |
| **Fig.2G** | WT : n=10<br>VGLUT3$^{T8I/T8I}$ : n=7-9 | Max z-scored Δ F/F (ACh) | | | | |
| | | Mixed-effects model (REML) | Genotype | F(1,18) = 3.065 | =0.0970 | Sidak’s multiple comparisons test |
| | | | Stress | F(1,14) = 13.75 | =0.0023 | |
| | | | Interaction | F(1,14) = 3.322 | =0.0898 | |
| **Fig.2I** | WT : n=4<br>VGLUT3$^{T8I/T8I}$ : n=6 | AUC DA Attack | | | | |
| | | Mixed-effects model (REML) | Genotype | F(1,16) = 0.03375 | =0.8565 | |
| | | | Stress | F(1,16) = 0.001034 | =0.9747 | |
| | | | Interaction | F(1,16) = 1.118 | =0.3061 | |
| **Fig.2J** | WT : n=4<br>VGLUT3$^{T8I/T8I}$ : n=6 | Max z-scored Δ F/F (DA) | | | | |
| | | Mixed-effects model (REML) | Genotype | F(1,16) = 0.2785 | =0.6049 | |
| | | | Stress | F (1, 16) = 5,843e -005 | =0.9940 | |
| | | | Interaction | F(1,16) = 0.8175 | =0.3793 | |
| **Fig.2K** | WT : n=10<br>VGLUT3$^{T8I/T8I}$ : n=9 | Correlation AUC and SP | | | | |
| | | Spearman correlation (WT, VGLUT3$^{T8I/T8I}$ ) | AUC ACh CSDS10<br>SP SIO post-CSDS | WT: Rho = 0.1151<br>VGLUT3$^{T8I/T8I}$: Rho = 0.3167 | =0.2353<br>=0.1530 | |
| **Fig.3B** | WT : n=10-11<br>VGLUT3$^{T8I/T8I}$ : n=8-9 | AUC ACh Nose to nose (SI) | | | | |
| | | Mixed-effects model (REML) | Genotype | F(1,18) = 0.5855 | =0.4541 | Uncorrected Fisher’s LSD |
| | | | Stress | F(1,16) = 2.491 | =0.1340 | |
| | | | Interaction | F(1,16) = 4.982 | =0.0403 | |
| **Fig.3C** | WT : n=10-11<br>VGLUT3$^{T8I/T8I}$ : n=8-9 | Max z-scored Δ F/F (ACh) | | | | |
| | | Mixed-effects model (REML) | Genotype | F(1,18) = 0.3177 | =0.5799 | Sidak’s multiple comparisons test |
| | | | Stress | F(1,16) = 6.456 | =0.0218 | |
| | | | Interaction | F(1,16) = 1.830 | =0.1950 | |

| | | | | | |
|---|---|---|---|---|---|
| **Fig.3E** | WT : n=5<br>VGLUT3$^{T8I/T8I}$ : n=7 | AUC DA Nose to nose (SI) | | | |
| | | Two-way RM ANOVA | Interaction | F(1,10) = 3.511 | =0.0904 | |
| | | | Genotype | F(1,10) = 1.519 | =0.2460 | |
| | | | Stress | F(1,10) = 0.1874 | =0.6742 | |
| **Fig.3F** | WT : n=5<br>VGLUT3$^{T8I/T8I}$ : n=7 | Max z-scored Δ F/F (DA) | | | | |
| | | Two-way RM ANOVA | Interaction | F(1,10) = 2.528 | =0.1429 | |
| | | | Genotype | F(1,10) = 1.847 | =0.2040 | |
| | | | Stress | F(1,10) = 1.617 | =0.2323 | |
| **Fig.3G** | WT : n=9<br>VGLUT3$^{T8I/T8I}$ : n=8 | Correlation AUC and SP | | | | |
| | | Spearman correlation (WT, VGLUT3$^{T8I/T8I}$ ) | AUC ACh SI post-CSDS<br>SP SI post-CSDS | WT: Rho = -0.2333<br>VGLUT3$^{T8I/T8I}$: Rho = -0.7857 | =0.5457<br>=0.0208 | |
| **Fig.S1A** | WT : n=14<br>VGLUT3$^{T8I/T8I}$ : n=15 | Total distance traveled SI with CD1 | | | | |
| | | Two-way RM ANOVA | Interaction | F(1,27) = 1.427 | =0.2427 | Sidak's multiple comparisons test |
| | | | Genotype | F(1,27) = 2.893e-005 | =0.9957 | |
| | | | Stress | F(1,27) = 22.69 | <0.0001 | |
| **Fig.S1B** | WT : n=14<br>VGLUT3$^{T8I/T8I}$ : n=16 | Total distance traveled in the SI test with empty box | | | | |
| | | Two-way RM ANOVA | Interaction | F(1,28) = 2.069 | =0.1614 | |
| | | | Genotype | F(1,28) = 1.329 | =0.2588 | |
| | | | Stress | F(1,28) = 3.969 | =0.0562 | |
| **Fig.S1C** | WT : n=10<br>VGLUT3$^{T8I/T8I}$ : n=15 | Total distance traveled in the 3C test | | | | |
| | | Two-way RM ANOVA | Interaction | F(1,23) = 0.01213 | =0.9133 | Sidak's multiple comparisons test |
| | | | Genotype | F(1,23) = 0.1893 | =0.6675 | |
| | | | Stress | F(1,23) = 35.20 | <0.0001 | |
| **Fig.S1D** | WT : n=15<br>VGLUT3$^{T8I/T8I}$ : n=16 | Total distance traveled in the EZM | | | | |
| | | Two-way RM ANOVA | Interaction | F(1,29) = 1.467 | =0.2356 | Sidak's multiple comparisons test |
| | | | Genotype | F(1,29) = 0.5606 | =0.4600 | |
| | | | Stress | F(1,29) = 10.64 | =0.00228 | |

| | | | | | | |
|---|---|---|---|---|---|---|
| **Fig.S1E** | WT : n=9<br>VGLUT3$^{T8I/T8I}$ : n=11 | Time spent in light | | | | |
| | | Two-way RM ANOVA | Interaction | F(1,18) = 2.615 | =0.1233 | |
| | | | Genotype | F(1,18) = 0.3362 | =0.5692 | |
| | | | Stress | F(1,18) = 02.324 | =0.1448 | |
| **Fig.S1F** | WT : n=15<br>VGLUT3$^{T8I/T8I}$ : n=16 | Time spent grooming | | | | |
| | | Two-way RM ANOVA | Interaction | F(1,29) = 2.048 | =0.1631 | |
| | | | Genotype | F(1,29) = 0.001845 | =0.9660 | |
| | | | Stress | F(1,29) = 0.9792 | =0.3306 | |
| **Fig.S2A** | WT : n=15<br>VGLUT3$^{T8I/T8I}$ : n=17 | Interaction time | | | | |
| | | Two-way RM ANOVA | Interaction | F(1,30) = 0.9470 | =0.3383 | Sidak's multiple comparisons test |
| | | | Genotype | F(1,30) = 0.2363 | =0.6305 | |
| | | | Stress | F(1,30) = 21.00 | <0.0001 | |
| **Fig.S2B** | WT : n=15<br>VGLUT3$^{T8I/T8I}$ : n=17 | Social preference index (CD1) | | | | |
| | | Two-way RM ANOVA | Interaction | F(1,30) = 0.06286 | =0.8037 | Sidak's multiple comparisons test |
| | | | Genotype | F(1,30) = 0.06772 | =0.7965 | |
| | | | Stress | F(1,30) = 37.74 | <0.0001 | |
| **Fig.S2C** | WT : n=13<br>VGLUT3$^{T8I/T8I}$ : n=15 | Social preference index (B6) | | | | |
| | | Two-way RM ANOVA | Interaction | F(1,26) = 0.06603 | =0.7992 | |
| | | | Genotype | F(1,26) = 0.008858 | =0.9257 | |
| | | | Stress | F(1,26) = 0.1564 | =0.6957 | |
| **Fig.S2D** | WT : n=12<br>VGLUT3$^{T8I/T8I}$ : n=14 | Social preference (z-scored) | | | | |
| | | Two-way RM ANOVA | Interaction | F(1,24) = 0.05661 | =0.8139 | Sidak's multiple comparisons test |
| | | | Genotype | F(1,24) = 0.05223 | =0.8212 | |
| | | | Stress | F(1,24) = 40.32 | <0.0001 | |
| **Fig.S2E** | WT : n=12<br>VGLUT3$^{T8I/T8I}$ : n=14 | Delta social preferences (z-scored) | | | | |
| | | Unpaired t test two-tailed (WT vs. VGLUT3$^{T8I/T8I}$) | | t = 0.2379 et DF = 24 | =0.8139 | |

| | | | | | | |
|---|---|---|---|---|---|---|
| | | One sample t test (WT, $VGLUT3^{T8I/T8I}$) | | t = 4.757 et DF = 11<br><br>t = 4.303 et DF = 13 | =0.0006<br><br>=0.0009 | |
| **Fig.S2G-H** | WT : n=12<br><br>$VGLUT3^{T8I/T8I}$ : n=14 | Multiple logistic regression (dependent variable = genotype) | | | | |
| | | Odds ratios estimate | Intercept<br>SI<br>3C<br>SI :3C | =0.8231<br>=0.9976<br>=1.873<br>=1.064 | =0.7707<br>=0.9889<br>=0.5848<br>=0.8423 | |
| | | Hosmer-Lemeshow test | | | = 0.8483 | |
| | | Tjur's R squared | | =0.02478 | | |
| | | ROC Curve | | Area = 0.5476 | =0.6807 | |
| | | Negative predictive power (%) | | =52.38 | | |
| | | Positive predictive power (%) | | =40.00 | | |
| **Fig.S2I** | WT : n=16<br><br>$VGLUT3^{T8I/T8I}$ : n=18 | Times spent in open arms | | | | |
| | | Two-way RM ANOVA | Interaction | F(2,64) = 2.063 | =0.1355 | Sidak's multiple comparisons test |
| | | | Genotype | F(1,32) = 0.1159 | =0.7358 | |
| | | | Stress | F(1.831,58.61) = 48.67 | <0.0001 | |
| **Fig.S2J** | WT : n=13<br><br>$VGLUT3^{T8I/T8I}$ : n=15 | Time spent in light | | | | |
| | | Two-way RM ANOVA | Interaction | F(1,26) = 0.003827 | =0.9511 | |
| | | | Genotype | F(1,26) = 0.3164 | =0.5786 | |
| | | | Stress | F(1,26) = 2.206 | =0.1495 | |
| **Fig.S2K** | WT : n=13<br><br>$VGLUT3^{T8I/T8I}$ : n=14 | Time spent grooming | | | | |
| | | Two-way RM ANOVA | Interaction | F(1,25) = 0.8409 | =0.3679 | |
| | | | Genotype | F(1,25) = 0.03739 | =0.8482 | |
| | | | Stress | F(1,25) = 0.6396 | =0.4314 | |
| **Fig.S3A** | WT males: | Interaction time | | | | |

| | | | | | | |
|---|---|---|---|---|---|---|
| | n=15<br><br>WT females : n=15 | Two-way RM ANOVA | Interaction | F(1,28) = 0.08216 | =0.7765 | Uncorrected Fisher's LSD |
| | | | Sex | F(1,28) = 0.3465 | =0.5608 | |
| | | | Stress | F(1,28) = 14.55 | =0.0007 | |
| **Fig.S3B** | WT males: n=10<br><br>WT females : n=13 | Social preference index (B6) | | | | |
| | | Two-way RM ANOVA | Interaction | F(1,21) = 4.266 | =0.0514 | |
| | | | Sex | F(1,21) = 3.632 | =0.0704 | |
| | | | Stress | F(1,21) = 1.598 | =0.2200 | |
| **Fig.S3C** | WT males: n=15<br><br>WT females : n=16 | Time spent in open arms | | | | |
| | | Two-way RM ANOVA | Interaction | F(1,29) = 15.68 | =0.0004 | Uncorrected Fisher's LSD |
| | | | Sex | F(1,29) = 16.52 | =0.0003 | |
| | | | Stress | F(1,29) = 37.78 | <0.0001 | |
| **Fig.S3D** | WT males: n=15<br><br>WT females : n=16 | % decreased time in open arms | | | | |
| | | Unpaired t test two-tailed (WT males vs. WT females) | | t = 2.117 et DF = 29 | =0.0429 | |
| **Fig.S3E** | WT males: n=10<br><br>WT females : n=12 | Principal component analysis (PCA) | | | | |
| | | Eigenvalue | PC1 | 1.758 | | |
| | | | PC2 | 0.7299 | | |
| | | | PC3 | 0.5117 | | |
| | | Proportion of variance | PC1 | 58.61% | | |
| | | | PC2 | 24.33% | | |
| | | | PC3 | 17.06% | | |
| | | Loading values PC1 | SI | 0.7902 | | |
| | | | 3C | -0.6836 | | |
| | | | EZM | -0.8165 | | |
| | | Loading values PC2 | SI | 0.3904 | | |
| | | | 3C | 0.7246 | | |
| | | | EZM | -0.2289 | | |
| **Fig.S3G-H** | WT males: n=15 | Multiple logistic regression (dependant variable = sex) | | | | |

| | | | | | |
|---|---|---|---|---|---|
| | WT females : n=15 | Odds ratios estimate | Intercept | =0.09916 | =0.0743 |
| | | | EZM | =4.499 | =0.1168 |
| | | | SI | =0.4770 | =0.1391 |
| | | | EZM:SI | =1.430 | =0.3047 |
| | | Hosmer-Lemeshow test | | | = 0.9858 |
| | | Tjur's R squared | | =0.2294 | |
| | | ROC Curve | | Area = 0.7422 | =0.0238 |
| | | Negative predictive power (%) | | =72.73 | |
| | | Positive predictive power (%) | | =63.16 | |
| **Fig.S4B1** | WT : n=3<br>VGLUT3$^{T8I/T8I}$ : n=6 | Peak value CSDS1 | | | |
| | | Wilcoxon Signed Rank Test (WT, VGLUT3$^{T8I/T8I}$) | | | =0.2500<br>=0.0312 |
| **Fig.S4B2** | WT : n=3<br>VGLUT3$^{T8I/T8I}$ : n=6 | Peak lag CSDS1 | | | |
| | | Wilcoxon Signed Rank Test (WT, VGLUT3$^{T8I/T8I}$) | | | =0.5000<br>=0.0625 |
| **Fig.S4C1** | WT : n=3<br>VGLUT3$^{T8I/T8I}$ : n=6 | Peak value CSDS10 | | | |
| | | Wilcoxon Signed Rank Test (WT, VGLUT3$^{T8I/T8I}$) | | | =0.2500<br>=0.0625 |
| **Fig.S4C2** | WT : n=3<br>VGLUT3$^{T8I/T8I}$ : n=6 | Peak lag CSDS10 | | | |
| | | Wilcoxon Signed Rank Test (WT, VGLUT3$^{T8I/T8I}$) | | | >0.9999<br>=0.1875 |
| **Fig.S4D1** | WT : | Peak value SI pre-CSDS | | | |

| | | | | | |
|---|---|---|---|---|---|
| | n=4<br><br>$VGLUT3^{T8I/T8I}$ :<br>n=6 | Wilcoxon Signed Rank Test (WT, $VGLUT3^{T8I/T8I}$) | | | =0.1250<br>=0.0312 |
| **Fig.S4D2** | WT :<br>n=4<br><br>$VGLUT3^{T8I/T8I}$ :<br>n=6 | Peak lag SI pre-CSDS | | | |
| | | Wilcoxon Signed Rank Test (WT, $VGLUT3^{T8I/T8I}$) | | | =0.2500<br>=0.0312 |
| **Fig.S4E1** | WT :<br>n=4<br><br>$VGLUT3^{T8I/T8I}$ :<br>n=6 | Peak value SI post-CSDS | | | |
| | | Wilcoxon Signed Rank Test (WT, $VGLUT3^{T8I/T8I}$) | | | =0.2500<br>=0.0312 |
| **Fig.S4E2** | WT :<br>n=4<br><br>$VGLUT3^{T8I/T8I}$ :<br>n=6 | Peak value SI post-CSDS | | | |
| | | Wilcoxon Signed Rank Test (WT, $VGLUT3^{T8I/T8I}$) | | | =0.1250<br>=0.0312 |

#### SUPPLEMENTARY FIGURES

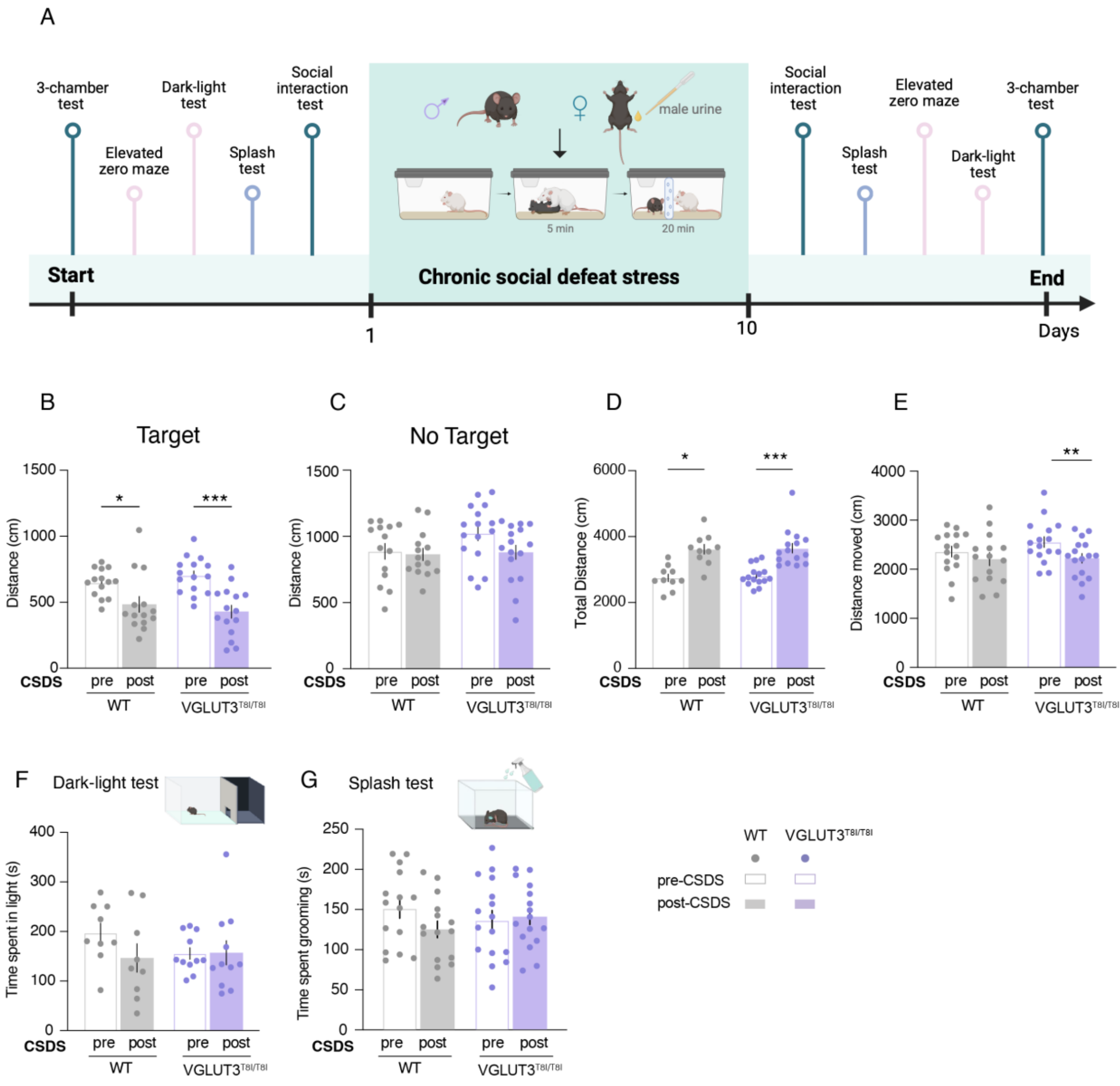

**Supplementary Figure 1. VGLUT3-p.T8I social phenotypes after CSDS are not confounded by general locomotor alterations**. **(A)** Experimental timeline. **(B)** Distance traveled in the target condition in the SI tests decreased after CSDS in both genotypes. **(C)** Distance traveled during the no-target condition in the social interaction (SI) tests showed no significant CSDS or genotype effect. **(D)** Total distance traveled during the three-chamber (3C) test increased after CSDS in both genotypes. **(E)** Total distance traveled in the elevated zero maze (EZM) decreased after CSDS in mutant mice but not in WT mice. **(F)** Time spent in the light compartment of the dark-light test revealed no CSDS or genotype effect. **(G)** Time spent grooming during the acute splash test pre- and post- CSDS showed no stress or genotype difference. WT mice, n=9-15 (grey bars); mutant mice, n=11-16 (purple bars); *$p<0.05$, **$p<0.01$, ***$p<0.001$(CSDS effect; Sidak's post-hoc, two-way RM ANOVA).

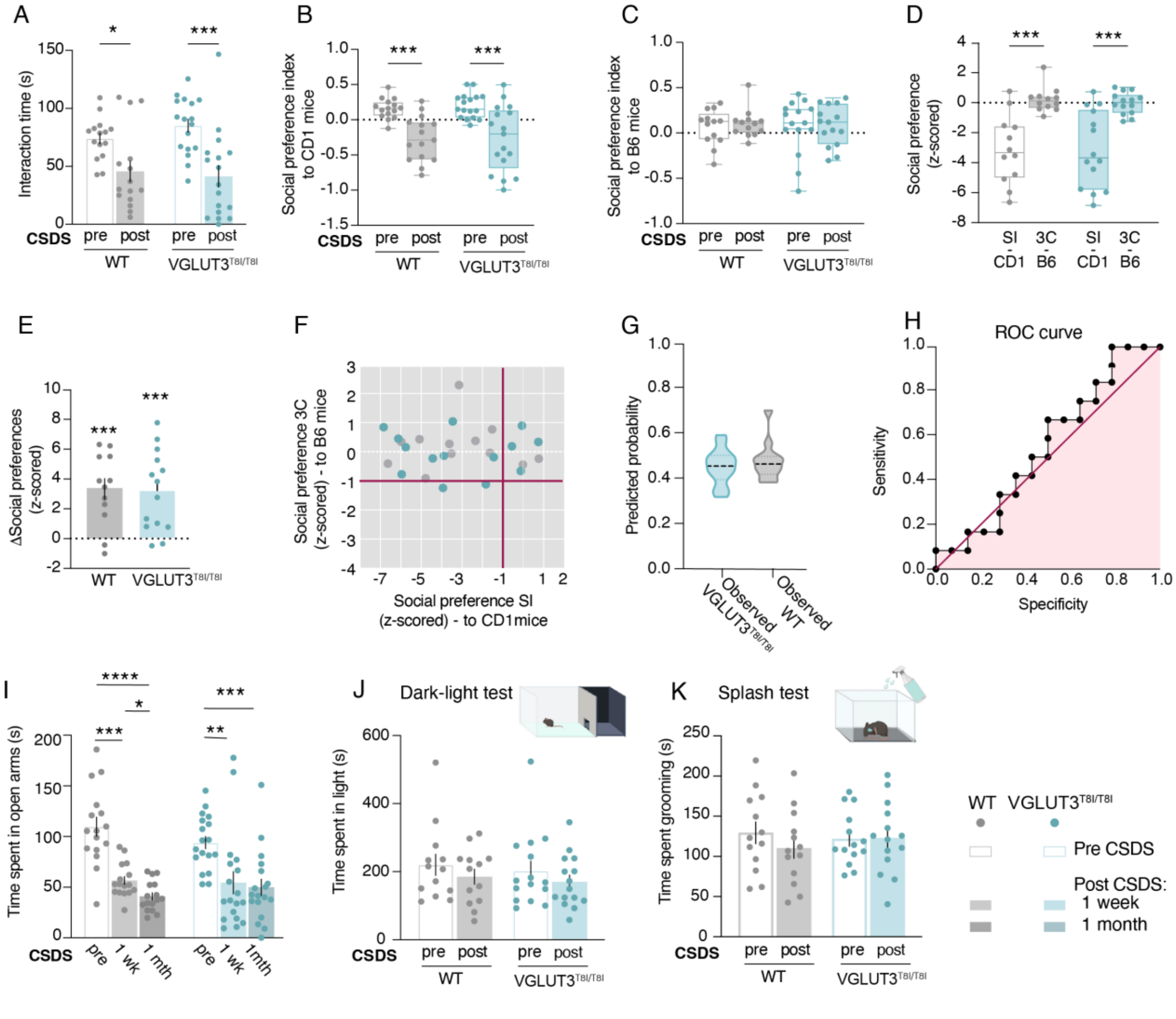

**Supplementary Figure 2. Female WT and VGLUT3$^{T8I/T8I}$ mice display a male-mutant-like phenotype following CSDS. (A)** Interaction time with the CD1 aggressor-strain mouse in the social interaction (SI) test decreased post-CSDS in both genotypes. **(B)** Social preference (SP) indices toward the CD1 aggressor strain decreased post-CSDS in both genotypes. **(C)** No significant CSDS or genotype effect on SP indices for a safe-associated B6 conspecific in the three-chamber (3C) test. **(D)** Z-scored SP values revealed selective CD1 avoidance in both genotypes. **(E)** Both genotypes showed greater ΔSP toward the CD1 aggressor strain than toward a safe-associated B6 conspecific. **(F)** Scatter plots of z-scored SP indices indicated similar social behavioral profiles between genotypes. **(G)** Logistic regression using B6 vs CD1 SP failed to separate WT from VGLUT3$^{T8I/T8I}$ females. **(H)** ROC curve confirmed poor genotype prediction from social behavior (AUC= 0.5476, p=0.6807). **(I)** Time spent in the open arms of the elevated zero maze (EZM) decreased post-CSDS for up to one month in both genotypes. **(J)** Time spent in the light compartment of the dark-light test showed no CSDS or genotype effect. **(K)** Time spent grooming in the acute splash test showed no CSDS or genotype effects. WT mice, n=12-16 (grey bars); mutant mice, n=14-18 (blue bars); *p<0.05, **p<0.01, ***p<0.001 (CSDS effect; Two-way RM ANOVA, Sidak post hoc comparisons; one sample t test).

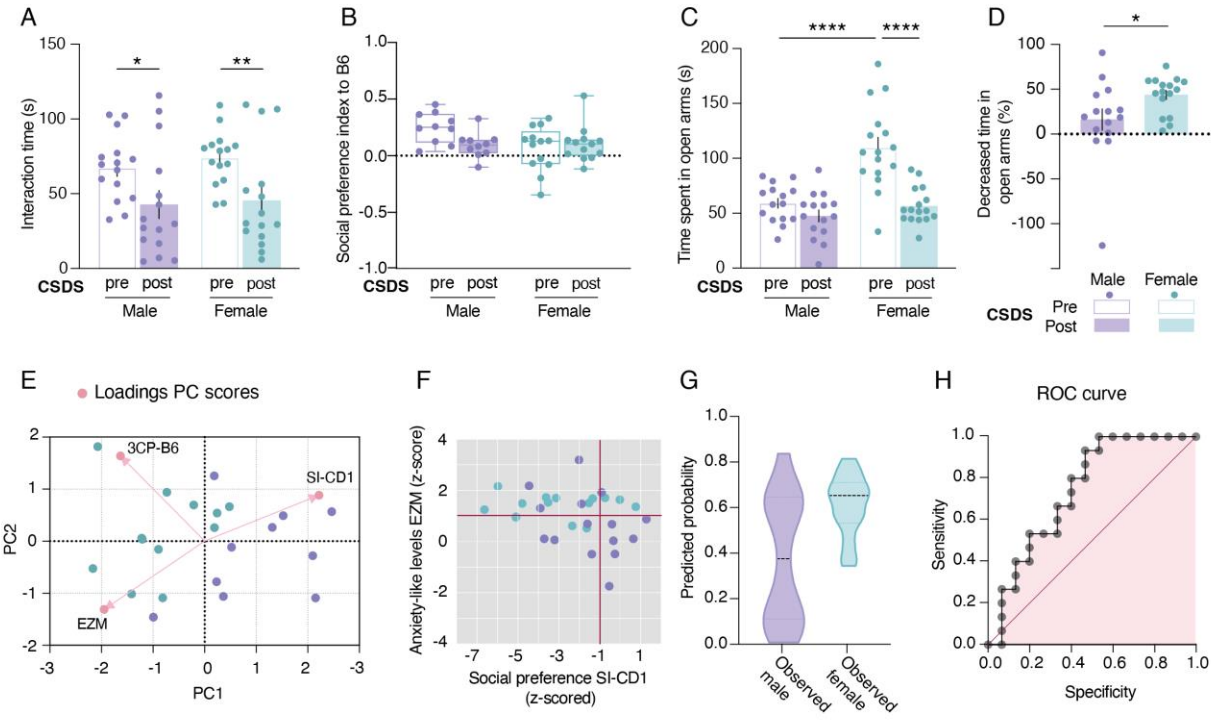

**Supplementary Figure 3. CSDS drives sex-dependent social avoidance and anxiety-like behavior in WT mice. (A)** Interaction time with the CD1 aggressor-strain mouse in the social interaction (SI) test decreased post-CSDS in both male and female WT mice. **(B)** Social preference (SP) indices for a B6 conspecific in the three-chamber (3C) test showed no significant CSDS or sex effect. **(C)** Time spent in open arms of the elevated zero maze (EZM) lower at baseline (pre-CSDS) in males compared to females and showed a post-CSDS decreased in females only. **(D)** Female WT mice exhibited greater CSDS-induced reduction in EZM open arm time than males. **(E)** Principal component analysis (PCA) scores plot with loading vectors revealed distinct sex-dependent behavioral clusters driven by anxiety-like behavior (EZM) and SP toward the CD1 aggressor strain along PC1 and PC2. **(F)** Scatter plots of z-scored SP indices revealed clear sex separation. **(G)** Logistic regression using CD1 SP and EZM robustly classified male versus female WT mice. **(H)** ROC curve confirmed strong model discriminative power (AUC=0.7422, p=0.0238). WT female mice, n=12-16 (blue bars); WT male mice, n=10-15 (purple bars); *p<0.05, ****p<0.0001 (CSDS effect), $^{\#\#}$p<0.01, $^{\#\#\#\#}$p<0.0001 (sex effect; two-way RM ANOVA, Sidak post hoc comparisons; unpaired t-test).

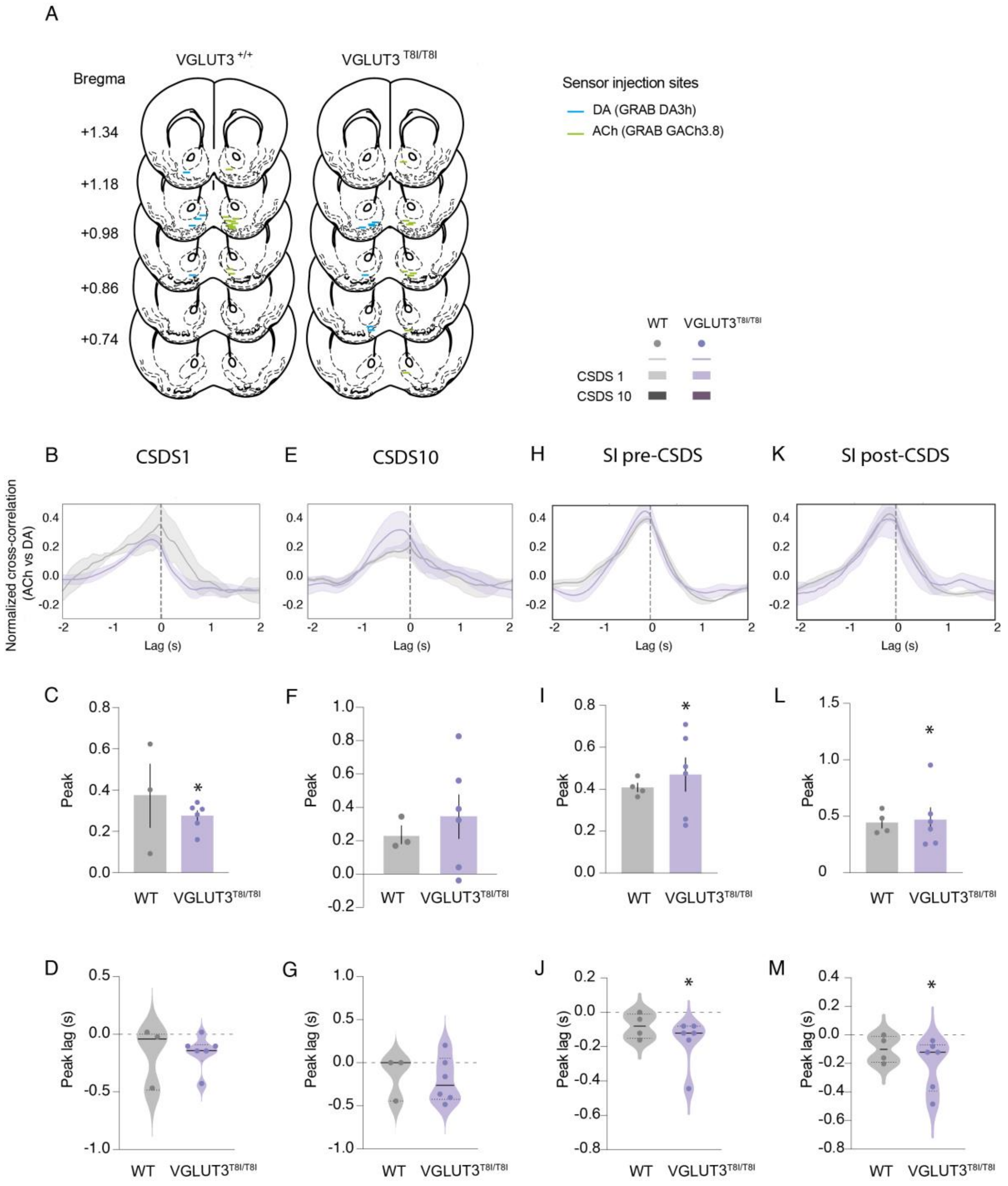

**Supplementary Figure 4. Histological verification of fiber photometry injection sites, viral expression and optic fiber placements. (A)** Schematic coronal sections illustrating individual optic fiber tip placement relative to bregma across anteroposterior (AP) levels for WT and VGLUT3$^{T8I/T8I}$ male mice. Inclusion required accurate nucleus accumbens medial shell (NAcMsh) targeting of at least one sensor (GACh3.8 and/or GRAB-DA3h). **(B)** ACh/DA cross-correlations on CSDS1 showed **(C)** comparable peak magnitude across genotypes and **(D)** peaks near 0 lag. **(E)** ACh/DA cross-correlations on CSDS day 10 showed **(F)** no peak magnitude difference between genotypes and **(G)** that temporal coupling between ACh and DA signals persisted without genotype-specific alterations. **(H)** ACh/DA cross-correlations during SI pre-CSDS showed **(I)** strong peak magnitudes for both genotypes and showed **(J)** a significant negative peak lag in mutant mice only suggesting that DA preceded ACh in mutant mice. **(K)** ACh/DA cross-correlations during SI post-CSDS showed **(L)** equivalent magnitudes across genotypes, **(M)** with persistent DA precedence over ACh in mutants.
